# Confirmation bias optimizes reward learning

**DOI:** 10.1101/2021.02.27.433214

**Authors:** Tor Tarantola, Tomas Folke, Annika Boldt, Omar D. Pérez, Benedetto De Martino

## Abstract

Confirmation bias—the tendency to overweight information that matches prior beliefs or choices—has been shown to manifest even in simple reinforcement learning. In line with recent work, we find that participants learned significantly more from choice-confirming outcomes in a reward-learning task. What is less clear is whether asymmetric learning rates somehow benefit the learner. Here, we combine data from human participants and artificial agents to examine how confirmation-biased learning might improve performance by counteracting decisional and environmental noise. We evaluate one potential mechanism for such noise reduction: visual attention—a demonstrated driver of both value-based choice and predictive learning. Surprisingly, visual attention showed the opposite pattern to confirmation bias, as participants were most likely to fixate on “missed opportunities”, slightly dampening the effects of the confirmation bias we observed. Several million simulated experiments with artificial agents showed this bias to be a reward-maximizing strategy compared to several alternatives, but only if disconfirming feedback is not completely ignored—a condition that visual attention may help to enforce.

## INTRODUCTION

Thriving in complex environments requires efficiently learning from the consequences of our actions. By this standard, decision making often appears persistently suboptimal due to various biases. Especially pervasive is confirmation bias, a tendency to learn from new information in a way that confirms prior beliefs (Nickerson, 1998). Confirmation bias is thought to explain an array of socially significant phenomena, including political polarization (Kaplan et al., 2016; Lord et al., 1979; Nyhan and Reifler, 2010; Taber and Lodge, 2006), radicalization (Rollwage et al., 2018), and the underestimation of risks posed by threats such as climate change (Kahan et al., 2012).

Recently, several studies have indicated that confirmation bias may influence even basic cognitive functions (Bronfman et al., 2015; Fontanesi et al., 2019; Palminteri et al., 2017; Urai et al., 2019), including processes as fundamental as reinforcement learning (Fontanesi et al., 2019; Palminteri et al., 2017). Participants have shown higher learning rates for outcomes that reinforce a choice—outcomes reflecting either that the choice was rewarded or that the unchosen option would have gone unrewarded if selected. This work suggests that confirmation bias, famous for its social and political significance, may in fact have its roots in the basic mechanics of reinforcement learning (Palminteri et al., 2017).

One largely unexplored possibility is that confirmation bias, like social-projection bias (Tarantola et al., 2017), benefits decision making by counteracting the effects of noise in both the environment (options with high reward rates sometimes yield neutral or bad outcomes) and the agent’s choice process (neural noise may cause less desirable choices). We interrogated this possibility by testing two related hypotheses. First, given the suggested noise-reducing role of visual attention during value-based choice (Krajbich & Rangel, 2012; Krajbich et al., 2010, 2011; Towal et al., 2013), we tested whether confirmation-biased reward learning stems from biased attention to choice-confirming feedback. And second, we tested whether confirmation bias leads to better performance during reward learning compared to other strategies (see Qiu et al., 2020; Rollwage et al., 2020). To address these questions, we collected eye-tracking and choice data from participants during a reinforcement learning task. We then compared performance data from different populations of artificial agents programmed with various combinations of choice- and outcome-contingent learning rates.

In line with recent work (Palminteri et al., 2017), participants showed substantially higher learning rates for choice-confirming outcomes than for choice-disconfirming outcomes. However, these differences were not driven by differences in visual attention. Rather, participants fixated the most on outcomes indicating missed opportunities— rewards accruing to unchosen options on trials where the chosen option went unrewarded. While it drew the most attention, this was the feedback from which participants learned the least. But despite this stark difference, we nevertheless found a positive main effect of fixation times on learning rates. In other words, more time spent looking at the outcome associated with a specific choice option on a given trial predicted higher learning rates for that choice option. In this way, visual attention may serve to dampen the opposing effects of an independently acting confirmation bias.

In addition, artificial agents that exhibited confirmation bias earned significantly more rewards on average than unbiased or differently biased actors. But agents that completely ignored choice-disconfirming outcomes did not fare as well. These results show that certain levels of confirmation bias—but not total blindness to disconfirming evidence—can help optimize reward learning in noisy environments. When, as in the real world, choices and rewards are stochastic, using previous choices as a kind of evidence may improve performance by counteracting some of this noise. These results may also shed light on the benefits and dangers of confirmation bias in more complex contexts, such as the consumption of false news stories and the perpetuation of filter bubbles (Knobloch-Westerwick, Mothes, & Polavin, 2020; Spohr, 2017).

## RESULTS

### Confirmation Bias in Learning Rates

We first tested whether participants would exhibit the same confirmation bias in reinforcement learning that others have observed (Palminteri et al., 2017). Thirty participants underwent eye tracking while completing a standard two-armed bandit reinforcement learning task. One participant was excluded for failing to learn (see Methods). Participants were asked to choose between two symbols, each with an independent probability of yielding a fixed monetary reward. After each choice, either an orange or a purple box was displayed around each symbol for three seconds (Fig. 1). The color of the box indicated whether a reward was (or, in the case of the unchosen symbol, would have been) earned.

**Figure 1.**
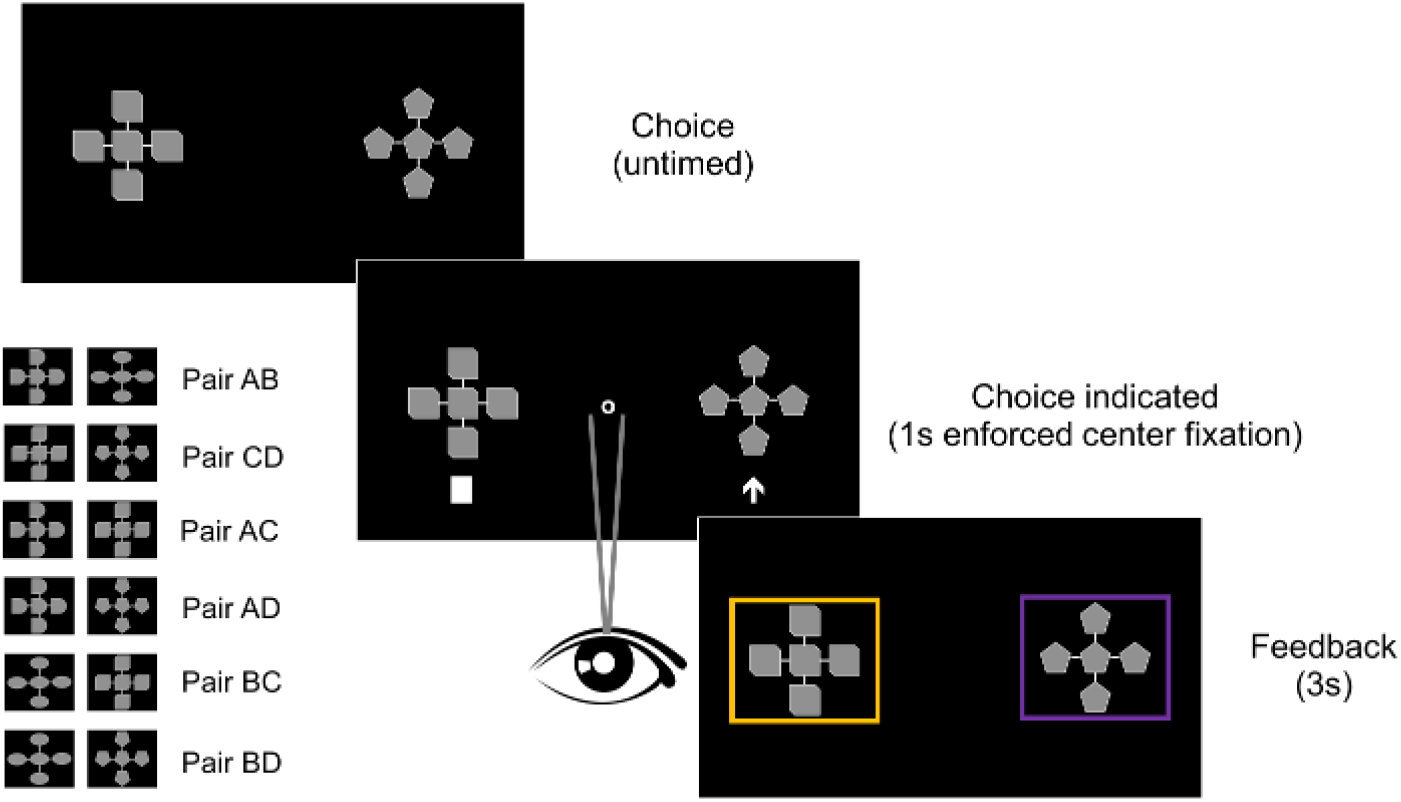
Learning task. Participants were asked to choose between one of two symbols on each trial, each with their own independent probability of yielding a reward. Once a choice was made, outcomes were displayed for both symbols after the participant fixated on a center marker for one second. Because the symbols’ reward probabilities were independent, sometimes both options yielded a reward, sometimes only one did, and sometimes neither did. The color of the box indicated the outcome (or, for the unchosen symbol, the foregone outcome). The color assigned to each outcome was counterbalanced across participants. Four different symbols were arranged in six unique pairs, and each pair was presented 40 times for a total of 240 trials. Each pair was roughly evenly spaced and left-right counterbalanced within blocks of 12 trials (see Methods).

Because participants were shown the outcomes of both options they had the opportunity to learn the value of each symbol regardless of their choice. Thus the task structure allows for the possibility of one fixed learning rate independent of choices and outcomes, and makes exploration unnecessary, allowing participants to focus exclusively on reward maximisation. Each symbol’s actual reward probabilities were independent and changed throughout the course of the task according to a random walk (Fig. 2). Across all trials, participants chose the better option—the option with the higher reward probability—three quarters of the time (mean=0.75; SEM=0.005, clustered by participant). Participants’ choices were rewarded an average of 62 percent of the time (mean=0.62; SEM=0.006, clustered by participant). We noted that participants tended to spend a substantial portion of the three-second feedback period fixating on the center of the screen rather than the outcomes (see Fig. 5). This reduced the risk of a ceiling effect, since fixations on outcomes were less likely to be cut short by the end of the feedback period.

**Figure 2.**
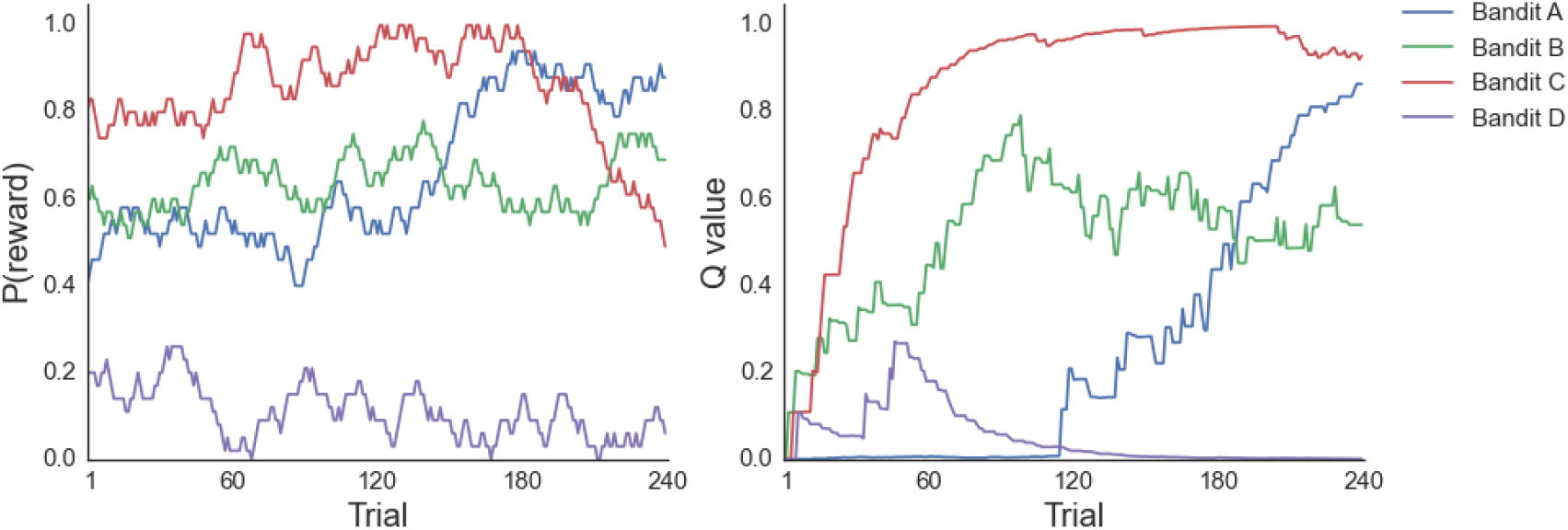
Actual reward probabilities (left) and modeled Q values (right) for Participant 25, shown as an example. Actual reward probabilities for the four symbols were initialized at 0.2, 0.4, 0.6, and 0.8, assigned randomly to different symbols. Each time the symbol was displayed during a trial, its actual reward probability increased or decreased randomly by 0.03, but was constrained within 0 and 1. The right-hand plot shows each trial’s mean Q value estimates from Model 6. All Q values were initialized at 0.

We modeled participants’ learning using a standard reward prediction-error reinforcement learning algorithm (Bush and Mosteller, 1951; Wakins and Dayan, 1992; Rescorla and Wagner, 1972) in which participants independently update the expected value of each symbol based on previously observed feedback. We compared five models with different learning-rate functions (Fig. 3). Model 1 used a fixed learning rate, and Models 2 through 4 allowed a symbol’s learning rate on a particular trial to vary based on whether the symbol was rewarded (Model 2), chosen (Model 3), or an additive combination of both (Model 4). Model 5, the confirmation-bias model, allowed learning rates to vary based on the confirmatory nature of the feedback. (See Methods for model specifications.) An outcome was *confirmatory* when it showed a reward for a chosen option or no reward for an unchosen option. An outcome was *non-confirmatory* when it showed no reward for a chosen option or a reward for an unchosen option. Model 5 was the most predictive (ELPD=-2550.5, SE=47.2) compared to the second-best, Model 4 (ELPD=-2596.7, SE=47.0; difference=46.2, SE=18.2; see Methods for model-fitting and model-comparison procedures).

**Figure 3.**
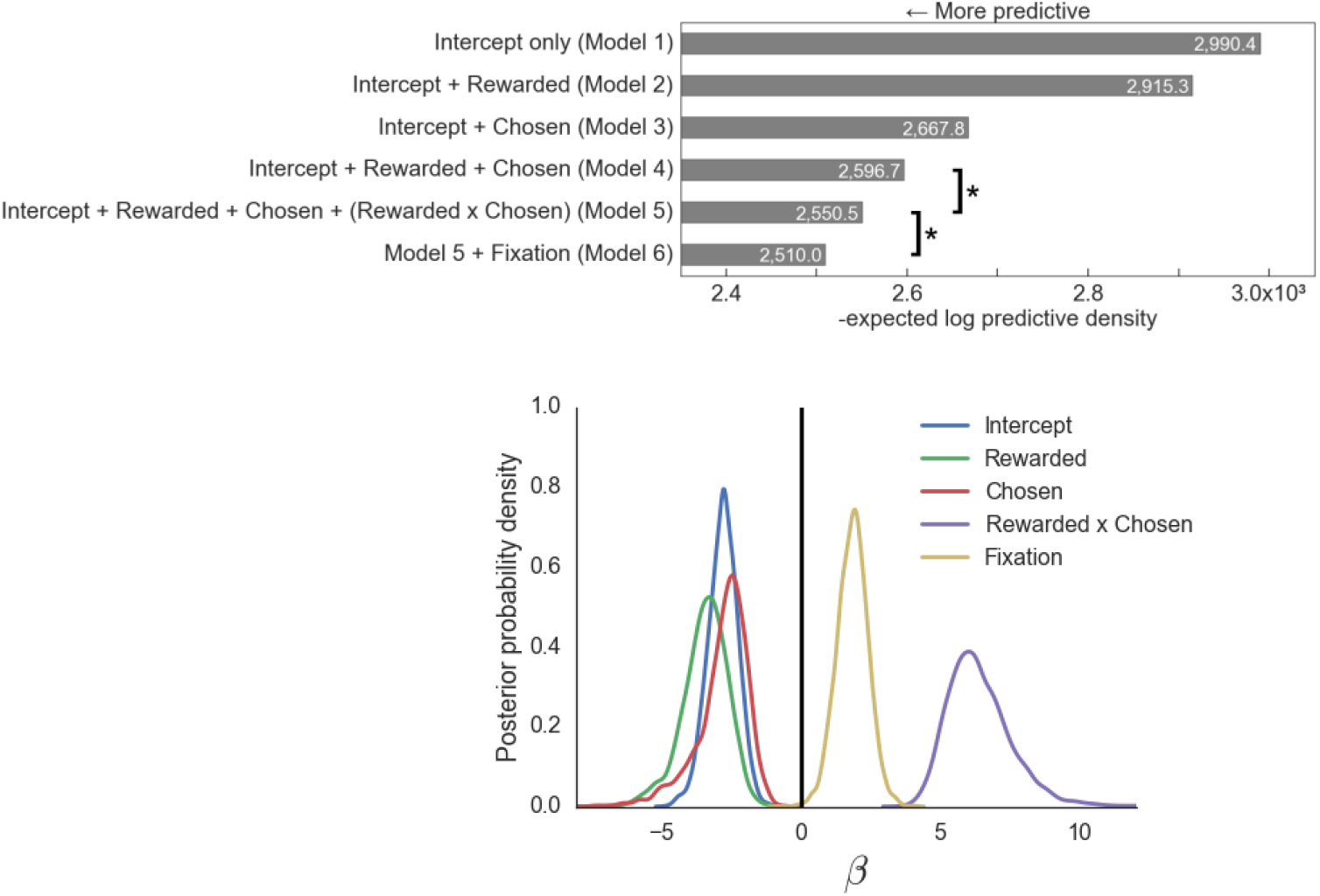
Learning model comparisons and parameter estimates. **(top)** Model 5 was significantly more predictive than the runner-up, as measured by the expected log predictive density for data from a new experiment. Model 6 was significantly more predictive than Model 5. *p<0.05, two-tailed. **(bottom)** Distributions indicate the uncertainty around the group mean parameter estimates for Model 6 and are kernel density smoothed.

To test whether learning rates were sensitive to visual attention, we then added outcome fixation times to Model 5. This model (Model 6) was significantly more predictive than Model 5 (ELPD=-2510.0, SE=47.1; difference=40.4, SE=10.4; see Methods for model specification). The parameters estimated from Model 6 yielded two main findings. First, consistent with previously published work (Palminteri et al., 2017), learning rates were highest for confirmatory outcomes—that is, for chosen symbols when they were rewarded (mean=0.14, SE=0.03 clustered by participant) and for unchosen symbols when they were not rewarded (mean=0.16, SE=0.03 clustered; Fig. 4, left). Learning rates were much lower for rewarded unchosen symbols (mean=0.05, SE=0.03 clustered) and unrewarded chosen symbols (mean=0.05, SE=0.02 clustered).

**Figure 4.**
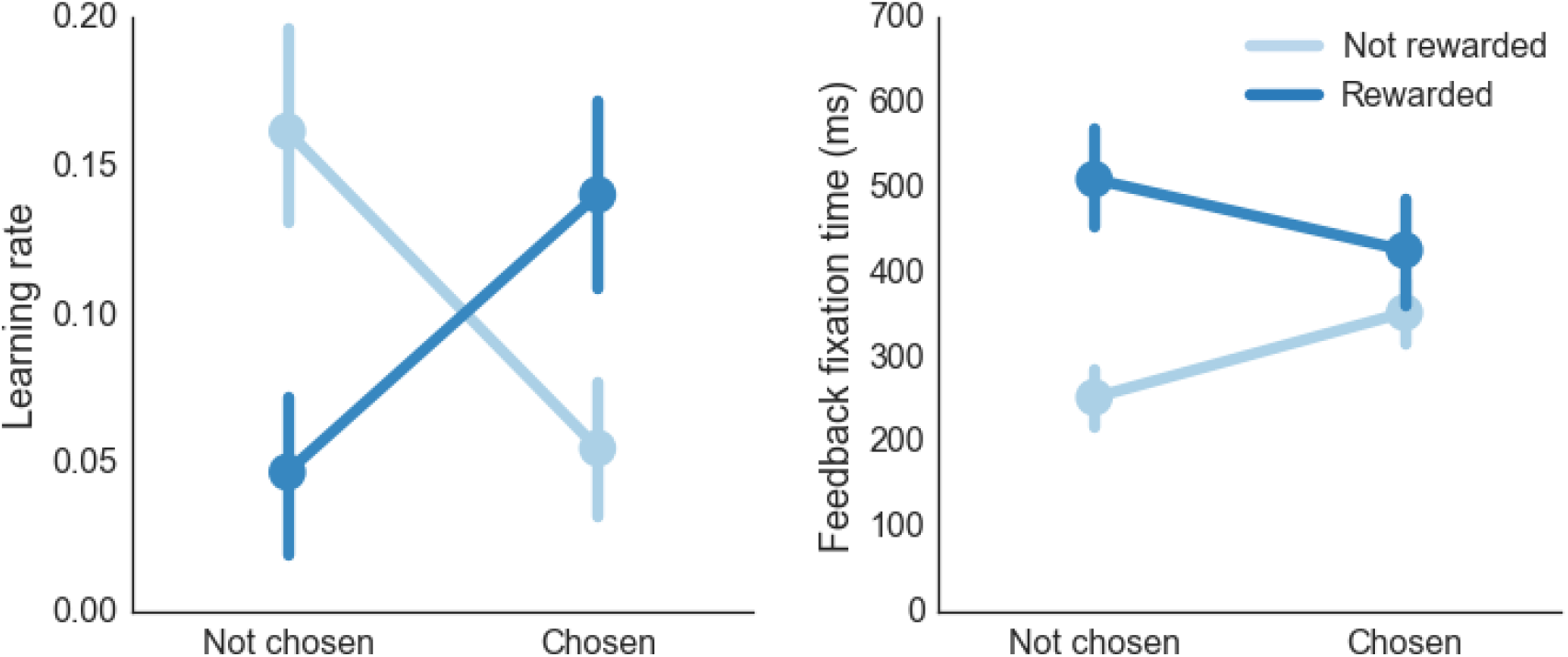
Learning rates and fixation times during feedback. **(left)** Learning rates were highest for chosen symbols when they were rewarded and for unchosen symbols when they were not rewarded. **(right)** By contrast, mean fixation times during the feedback phase were highest for unchosen symbols when they were rewarded and lowest for unchosen symbols when they were not rewarded. Learning rates are based on the mean estimates from Model 6 for each trial. Error bars indicate bootstrapped standard errors clustered by participant. Fixation data reflect 6,894 trials, which exclude 66 trials due to recording errors; these missing values were interpolated for the learning model.

Second, the time spent fixating on a symbol during the feedback period had a positive effect on learning rates, as indicated by the group mean parameter estimate (group mean estimate=1.86, SE=0.56). In other words, fixating on a symbol’s outcome predicted greater learning from that outcome.

### Fixation patterns

Fixation times themselves showed a strikingly different pattern from participants’ learning rates. When they were shown feedback, participants fixated most on rewarded unchosen symbols (mean=508ms, SE=58 clustered by participant) and fixated least on unchosen symbols that were not rewarded (mean=252 ms, SE=35 clustered; Fig. 4, right). In other words, despite a positive effect of fixation time on learning rates, fixations themselves seemed to concentrate most on the outcomes from which participants learned the least.

Closer inspection revealed that the difference observed in fixation times was driven by trials in which the outcomes for the two items differed. Participants were more likely to look at a symbol’s outcome if it was rewarded and the other symbol was not (Fig. 5, top panels). (Because each symbol’s outcome was independent, there were many trials in which neither symbol [23%] or both symbols [24%] were rewarded; see Fig. 5, bottom panels.) Participants were also more likely to fixate on at least one item during feedback if both outcomes disconfirmed their choice—that is, when the chosen symbol was unrewarded and the unchosen symbol was rewarded (a “missed opportunity” trial; Fig. 5, top right). We ran a multi-level logistic regression analysis predicting whether a participant fixated on a symbol’s outcome, treating each participant as a random effect with respect to the intercept. This analysis showed significant positive fixed effects of reward difference (−1 or 1) between an item and the other item on the screen (*β*=0.35, SE=0.02, z=14.5, p<10^-15^) and whether feedback indicated a missed opportunity *β* =1.51, SE=0.06, z=23.3, p<10^-15^). There was also a small positive effect of the absolute value of the Model 6 prediction error for that symbol *β* =0.26, SE=0.06, z=4.2, p<10^-4^), suggesting that the magnitude of surprise can independently draw attention to an outcome. Collectively, these results suggest that participants paid most attention to trials where the outcome between the two options differed. In these situations, visual attention shows a disconfirmation bias, the opposite pattern from what we see in the learning rates.

**Figure 5.**
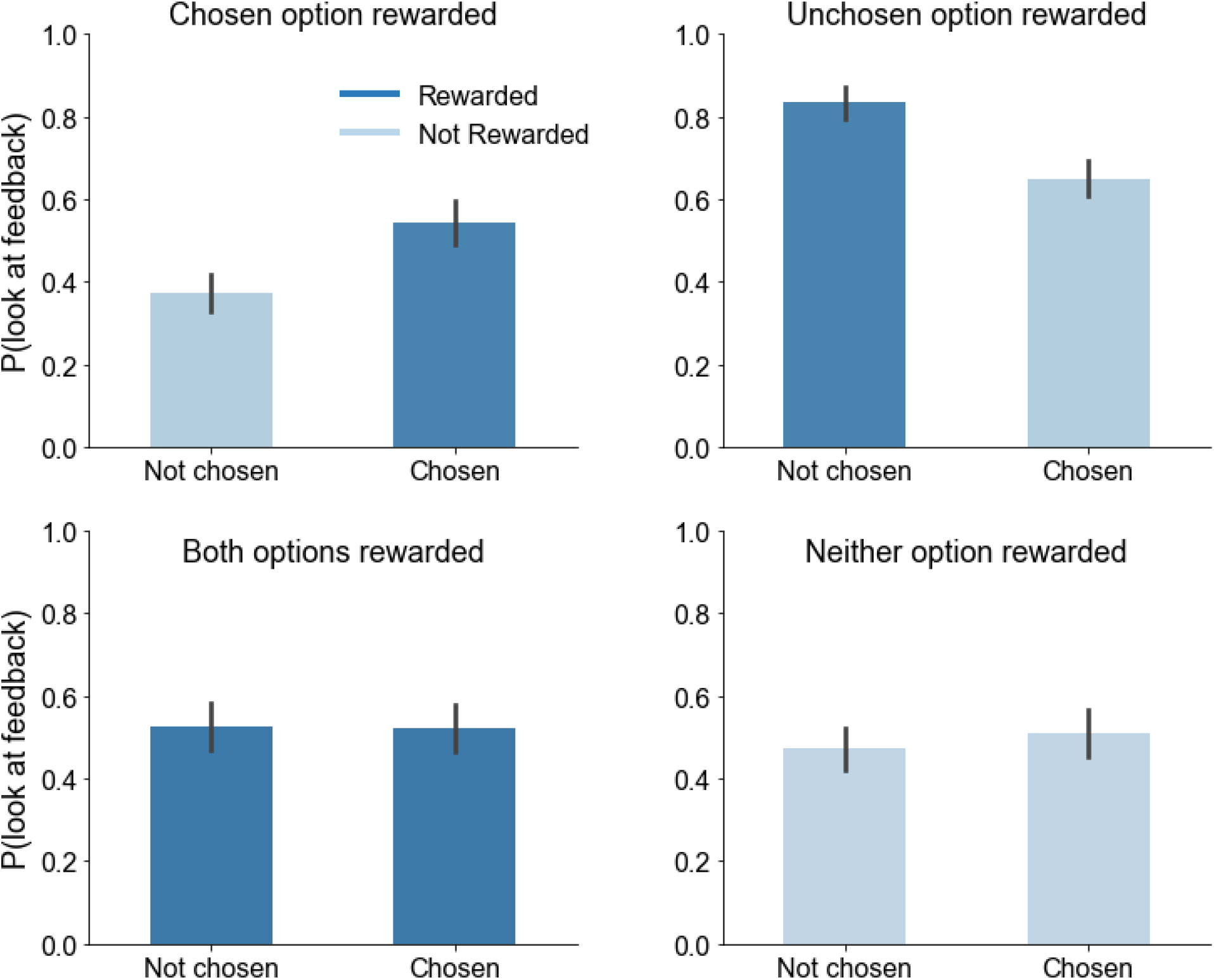
Probability of fixating on a symbol when its outcome was presented. Participants were more likely to look at a symbol’s outcome when it was rewarded and the other symbol presented was not (top panels, top left 38% of trials, top right: 15% of trials), compared to trials in which rewards accrued to both symbols (bottom left: 24% of trials) or neither symbol (bottom right: 23% of trials). In addition to this effect, participants were more likely to look at either symbol’s outcome in missed-opportunity trials, in which the chosen item was unrewarded and the unchosen item was rewarded (top right). Error bars indicate bootstrapped standard errors clustered by participant. Fixation data reflect 6,894 trials, which exclude 66 trials due to recording errors.

**Figure 6.**
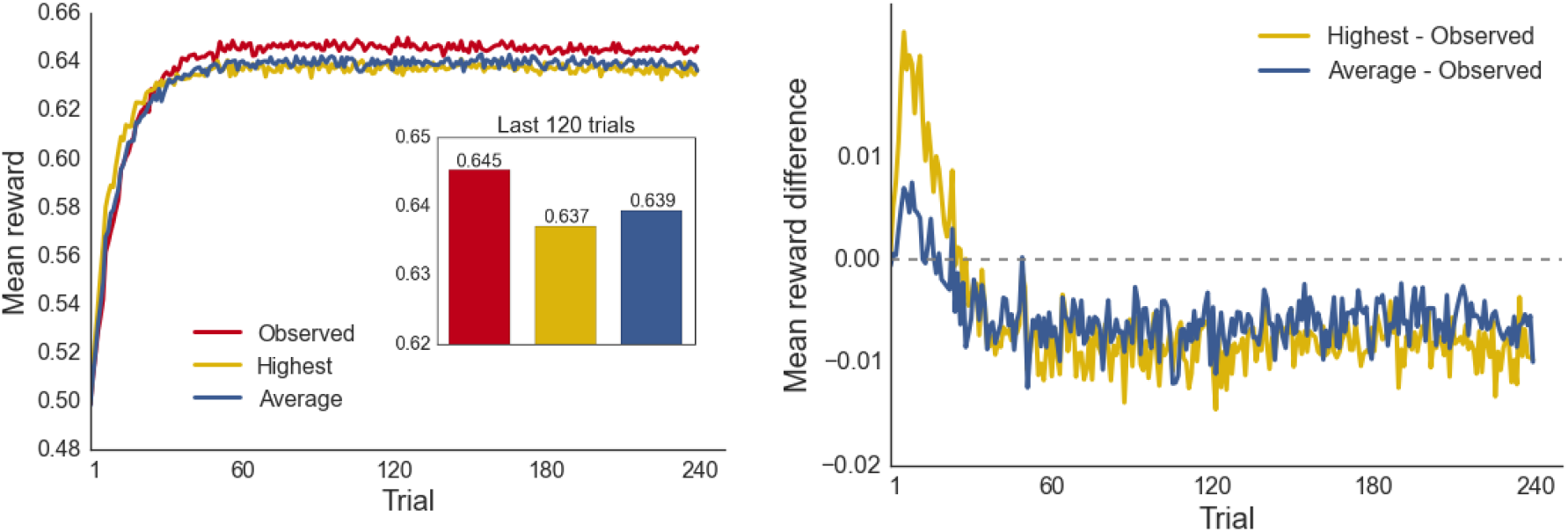
Performance of artificial agents using different learning strategies. **(left)** Mean rewards per trial from agents using the choice- and reward-dependent learning rates observed in our participants (red) compared to agents applying single learning rates equally to all outcomes (α = 0.16, yellow; α *=* 0.11, blue). Performance plateaued at a higher level for the observed compared to the other two strategies (inset shows means for the last 120 trials). **(right)** The mean reward from the *highest* (yellow) and *average* (blue) minus the *observed* strategy. While the *observed* strategy disadvantaged performance at the beginning of learning, it quickly led to a sustained advantage. Each strategy was simulated 100,000 times.

We also examined how long participants spent fixating on an outcome in the 52 percent of trials in which they fixated on either outcome at all. Fixation times in these cases showed a similar pattern (see Supplementary Materials).

These sets of fixation results stand in stark contrast to the learning rates revealed by our most predictive choice model. While participants’ attention was drawn most to feedback suggesting they made a mistake, this was the feedback that updated learned values the least.

### Effects on performance

As in recent studies, our results showed participants to be strikingly asymmetrical in how they learned from the outcomes of their choices. They learned roughly three times as much from outcomes that seemed to confirm their choices—rewards for chosen symbols and non-rewards for unchosen symbols—than from outcomes that did not. Is this a reward-maximizing strategy, or does it lead to worse performance?

To answer this question, we ran a series of simulated experiments and compared the average number of rewards earned by artificial agents programmed to exhibit one of three different learning strategies. In the first, *observed* strategy, agents adopted the learning strategy observed in our human participants. We programmed the agents to use the four mean learning rates observed in each of the choice-reward conditions (chosen rewarded=0.14, chosen unrewarded=0.05, unchosen rewarded=0.05, unchosen unrewarded=0.16) and apply them in the same way during our simulated experiments. In the second, *highest learning* strategy, agents applied the highest of these observed learning rates (0.16) equally to all choice-reward conditions. In the third, *average learning* strategy, agents used the average observed learning rate (0.11) and applied it equally to all choice-reward conditions. The softmax inverse temperature for all agents was set to 8.07, the mean of the group mean parameter estimate recovered from our human participants using Model 6. This ensured that all agents exhibited the same amount of stochasticity in converting learned Q-value differences into choices.

We first ran 100,000 simulated experiments for each of the three agent populations and compared the mean number of rewards they earned over the course of learning. We found that the *observed* strategy, while disadvantaging agents’ performance at the beginning of learning compared to the alternative strategies, quickly led to a sustained performance advantage (Fig. 7), suggesting that the strategy we observed in our human participants may be tuned to maximize long-term reward rates.

**Figure 7.**
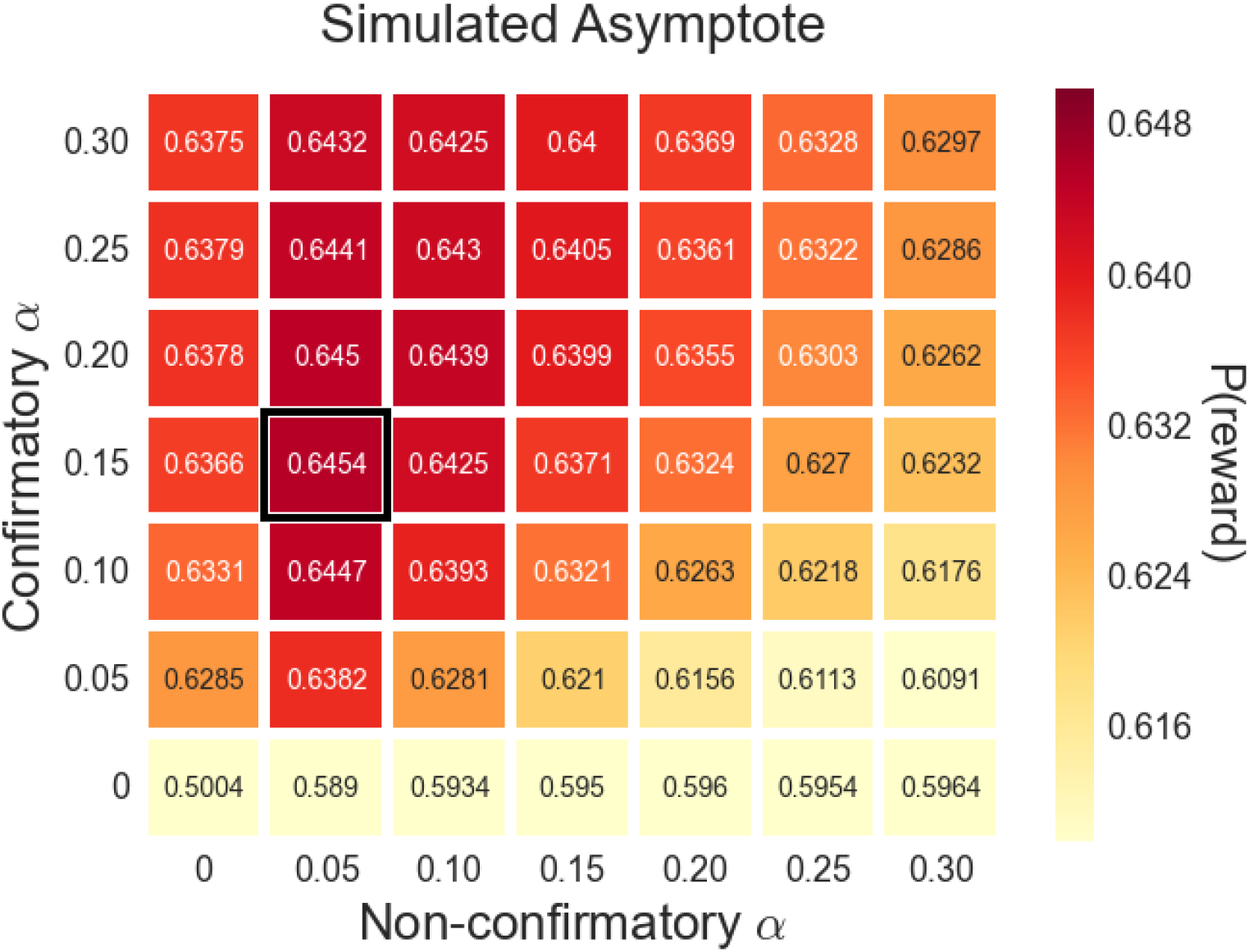
Mean sustained performance for different combinations of confirmatory and non-confirmatory learning rates. The observed strategy (black border) outperformed all others in the task. Each cell represents the mean reward outcome for 100,000 simulated participants in the last 120 trials.

We then compared the mean asymptotes—the average performance during the last 120 trials—of a number of different combinations of confirmatory (chosen rewarded/unchosen unrewarded) and non-confirmatory (chosen unrewarded/unchosen rewarded) learning rates (Fig. 8). We found that the *observed* strategy yielded the best sustained performance of any combination tested. We also noted a stark asymmetry along the diagonal, indicating that learning more from confirmatory than non-confirmatory outcomes—in other words, exhibiting a confirmation bias—provides clear advantages for many different learning-rate combinations.

**Figure 8.**
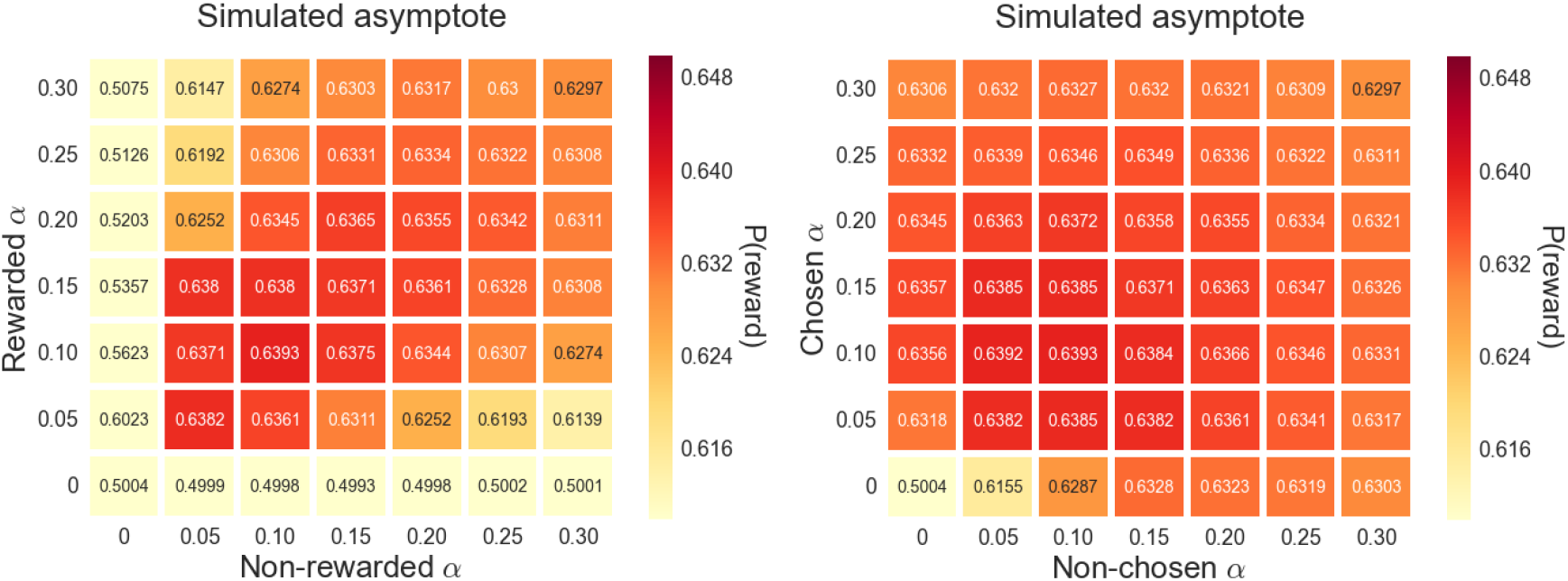
Mean sustained performance for different combinations of rewarded/non-rewarded learning rates (left) and chosen/non-chosen learning rates (right). In none of the combinations tested did performance exceed that of an unbiased learner with a simple learning rate of 0.10 for all outcomes. Consequently, none of these strategies reached the level of performance of many of the confirmatory/non-confirmatory strategies shown in Figure 8. Each cell represents the mean reward outcome for 100,000 artificial agents in the last 120 trials.

While the differences in reward probabilities appear small, we note that the total range of potential rewards was fairly constrained due to the design of the experiment. A test run of 100,000 simulated experiments showed that in only 67 percent of trials did the option with the higher actual reward probability yield a reward outcome. In other words, a perfect learner—an agent who chose the better option on every trial—would be rewarded only 67 percent of the time. And in 24 percent of trials, neither option was rewarded, meaning that even an omniscient agent would be rewarded only 76 percent of the time. By comparison, agents who chose at random (with learning rates set to 0) were rewarded 50 percent of the time.

While confirmation-biased agents showed clear performance advantages, these results also showed that completely ignoring non-confirmatory outcomes can hurt performance. Agents with non-confirmatory learning rates set to zero performed worse than many others, including some unbiased agent populations. This suggests that, while a confirmatory asymmetry in learning rates may be beneficial, learned values should still be somewhat sensitive to evidence that a previous choice was incorrect. The eye-gaze patterns we observed in our human participants may help to enforce this kind of open-mindedness, since visual fixations, which were drawn to choice-disconfirming evidence, also boosted learning rates for that evidence.

Lastly, we simulated the performance of two alternative types of learning strategies explored in previous literature: one in which learning rates differed depending on whether an item was rewarded or not, regardless of whether it was chosen (Cazé and Van Der Meer, 2013; Lefebvre et al., 2017; Palminteri et al., 2017; Perez and Dickinson, 2020); and one in which learning rates differed depending on whether an item was chosen or not, regardless of whether it was rewarded (Boorman et al., 2011; Li & Daw, 2011). In neither of these cases did any combination tested outperform a strategy using a single learning rate of 0.10 for all outcomes (Fig. 9). These strategies were even outperformed by several confirmatory/non-confirmatory strategies that performed worse than the strategy observed in our participants.

## DISCUSSION

Our results reveal a surprising dissociation between the drivers of learning and the drivers of visual attention during the feedback period. While visual attention toward an outcome was positively associated with increased learning from that outcome, it was not the primary driver of learning. Participants were most likely to look at outcomes during trials in which the chosen item was not rewarded and the unchosen item was—the outcomes from which they learned the least. Fixation probabilities and fixation times for an outcome were also predicted by the reward difference between that outcome and the other outcome presented on the screen. Participants’ eyes were drawn more to the rewarded outcome than the unrewarded one. When neither or both items were rewarded, there was no difference.

These effects suggest that visual attention in this context may serve chiefly to orient participants toward salient features in the visual field, including evidence that they made a mistake. This is supported by the additional positive effect we observed of prediction errors on fixation probabilities. In this way, given the effect of fixation time on learning rates, visual attention may serve to tamper the confirmation bias, helping to ensure that participants do not unduly ignore choice-disconfirming outcomes.

These results offer some new insight into the role of attention in learning more generally. To our knowledge, this is the first study to examine the relationship between learning and visual attention toward outcomes. Of course, attention in Pavlovian and instrumental learning has been a topic of inquiry for many decades (Beesley et al., 2015; Mackintosh, 1975; Pearce and Hall, 1980; Pearce and Mackintosh, 2010; Le Pelley, 2004; Le Pelley et al., 2011; Thrailkill et al., 2018). But this previous work has focused on the attention paid to stimuli that *predict* outcomes—and how human and non-human animals learn to discriminate important cues in the environment from irrelevant ones—rather than the attention paid to the outcomes themselves. Our results suggest that visual attention toward outcomes may help to store reward associations, much like fixations during value-based choice may help to retrieve them (see Cavanagh et al., 2014; Krajbich et al., 2010, 2012; Krajbich & Rangel, 2011; Towal et al., 2013). But our results also call into serious question the notion that visual attention during feedback is a primary driver of reinforcement learning. An apparently independent confirmation bias was much more impactful.

Results from our artificial agents revealed that this bias—being more sensitive to choice-confirming outcomes—can lead to a sustained performance advantage compared to neutral learning strategies. A key reason for this might be that exaggerating differences in learned values helps counteract the noise inherent in the choice process (Cazé and Van Der Meer, 2013; Tarantola et al., 2017; Tsetsos et al., 2016). Because choices are stochastic, the agent does not choose the option with the higher learned value one hundred percent of the time. Rather, the probability of choosing the option with the higher value increases as the difference in the values increases. For this reason, exaggerating this difference through a learning bias might result in better choices more often (Cazé and Van Der Meer, 2013; Tarantola et al., 2017). For this reason, biasing value updates in the direction of prior choices—which are correct more often than not—may help to reduce the incidence of stochastic errors. It may also help minimize the effect of external noise caused by the natural stochasticity in choice outcomes, allowing agents to downplay the effect of misleading feedback, such as the occasional foregone reward from a normally bad choice or a missed reward from a normally good choice. In essence, participants may be using their prior choices as useful evidence when updating learned values.

This explanation may also apply to the more complex yet fundamentally similar findings in the psychological literature on confirmation bias. Just as there is noise in the reward environment—sometimes a bad choice yields a good outcome, and vice versa— there might be noise in the social and political environment, where some information may be similarly unreliable. In the same way that we use prior choices as evidence to inform subsequent choices, we might use prior attitudes as evidence to inform future attitudes and insulate them against noise. Consequently, the type of political polarization observed in previous studies (Lord et al., 1979; Taber and Lodge, 2006) also resembles the polarization of *Q* values that leads to better performance (Cazé and Van Der Meer, 2013).

But our results also yield an important caveat. While they show a clear advantage of confirmation bias in reinforcement learning, they also show that choice-disconfirming outcomes should not be ignored altogether. Agents with non-confirmatory learning rates set to zero achieved worse asymptotic performance compared to many other learning-rate combinations, including the best-performing unbiased strategy. This makes intuitive sense—while treating prior choices as evidence can help us cut through noise, we should not be totally blind to evidence suggesting that our choices might have been wrong. The same could be said for decisions of social and political importance. While confirmation bias in these more complex contexts might still offer some advantages, total insulation from disconfirming evidence—caused by filter bubbles, for example—can hinder collective responses to significant public risks.

## METHODS

### Experimental procedure

Participants completed a two-armed bandit task with the aim to maximize their earnings. Each bandit symbol had the same reward magnitude (2.08 pence), but the underlying rate of reward differed. On each trial, two symbols were shown and participants made their choice with a key press. Following the choice, a circle appeared at the center of the screen to cue participants to fixate on the center before receiving feedback. Once participants fixated on the center circle for one second, the circle disappeared and boxes appeared around the symbols. The boxes were either orange or purple, the color indicating whether or not the bandit was rewarded. (The colors associated with reward and non-reward were counterbalanced across participants). The reward probabilities of the bandits were independent, so for any given trial both symbols could yield a reward, neither symbol could yield a reward, or one could yield a reward but not the other. To control for possible generalization effects, the symbols were chosen from a previous study that demonstrated little to no generalization between pairs of these stimuli (Pérez et al., 2018). Participants’ eye movements were recorded during all phases of each trial. Participants first completed 20 practice trials with two symbols that had stable reward rates (0.75 and 0.25) in order to familiarize themselves with the task structure and to make sure they understood the aim of the task. The practice trials did not count toward their earnings.

After the practice block, 240 main trials were administered. These trials had four different symbols, making six unique pairs that were each shown 40 times. The position of the bandits on the screen was counterbalanced for each pair within each 12-trial block. The four bandits started with reward rates of 0.2, 0.4, 0.6 and 0.8, assigned to random symbols at the beginning of the session. The reward rates for the main symbols were not stable throughout the experiment. Each time a symbol was shown, its reward rate changed by 0.03. If the reward rate of a bandit reached 0 or 1, it bounced to 0.03 or 0.97, but otherwise they were equally likely to increase or decrease. Participants were informed that the reward rates of the bandits would change over time.

### Participants

Thirty participants took part in the study. One participant was excluded for failing to meet our *a priori* inclusion criterion, which required that a logistic model including each trial’s difference in actual reward rates significantly predict each participant’s choices better than an intercept-only model. (This was not the case for the excluded participant, p=.70.) The final sample therefore included 29 participants (20 female) with a mean age of 26 (sd=6.37). All participants gave informed consent and were paid a £10 show-up fee and 2.08 pence per reward (a maximum of £5). The protocol was approved by the internal ethics committee of the Department of Psychology at the University of Cambridge.

### Eye tracking

Eye movements were recorded at 1,000Hz with an EyeLink 1000 Plus eye-tracker (SR-Research, Ontario). Interest areas were predefined as two rectangular areas (400 x 329 pixels) containing the symbols. These areas had the same dimensions and positions as the outcome boxes but were only visible when outcomes were presented.

### Computational modeling

We modeled participants’ learning using a simple *Q*-value update algorithm:

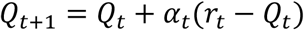

where *t* is the trial number for that particular symbol (equal to the number of times that particular symbol has been presented so far), *r* ∈ {0, 1} is a Kronecker delta indicating whether that symbol yielded a reward on trial *t*, and α is the learning rate. The learning rate indicates the extent to which a participant’s perceived expected value of a particular symbol, *Q*, is updated with each new outcome. A higher learning rate means that a participant updates her estimate of the symbol’s value to a greater degree based on a new piece of information. The learning rate also defines the magnitude with which participants weight recent feedback more heavily than less recent feedback.

We used a softmax choice rule to model each response as a function of differences in *Q* values:

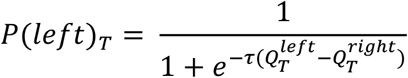

where the probability of a participant choosing the left symbol on experimental trial *T* is a logistic function of the difference in *Q* values at that trial, weighted by an inverse temperature parameter *τ*, which describes how sensitive choices are to *Q* value differences. A high inverse temperature means that even a small *Q* value advantage makes choosing a symbol much more probable, while a low inverse temperature means that the probability of choosing a symbol will be less affected by differences in *Q* values. In this way, τ operates as an index of noise in the choice process.

We tested five combinations of choice and reward effects on the learning rate, defining each learning rate for a particular symbol on trial *t* as a logistic function:

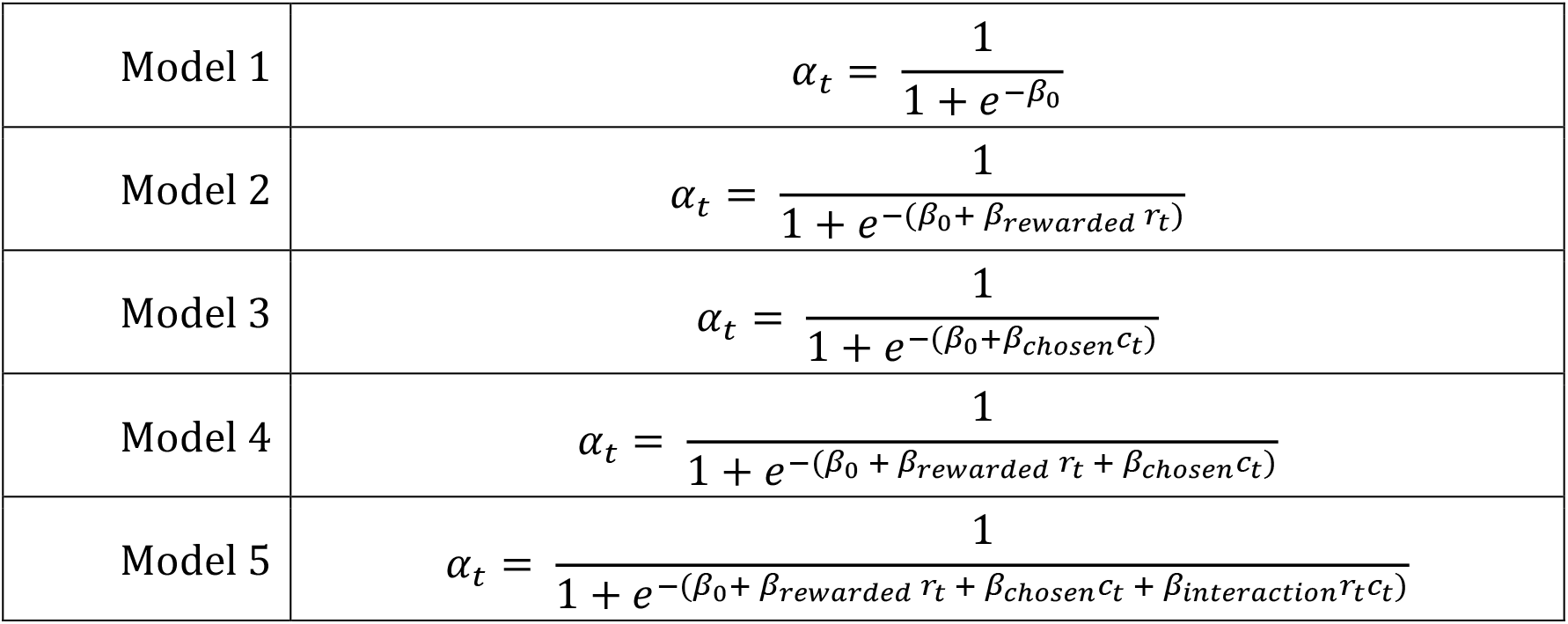

|  |  |
| --- | --- |
| Model 1 | $\alpha_t = \frac{1}{1 + e^{-\beta_0}}$ |
| Model 2 | $\alpha_t = \frac{1}{1 + e^{-(\beta_0 + \beta_{rewarded} r_t)}}$ |
| Model 3 | $\alpha_t = \frac{1}{1 + e^{-(\beta_0 + \beta_{chosen} c_t)}}$ |
| Model 4 | $\alpha_t = \frac{1}{1 + e^{-(\beta_0 + \beta_{rewarded} r_t + \beta_{chosen} c_t)}}$ |
| Model 5 | $\alpha_t = \frac{1}{1 + e^{-(\beta_0 + \beta_{rewarded} r_t + \beta_{chosen} c_t + \beta_{interaction} r_t c_t)}}$ |

where *c* and *r* are Kronecker deltas indicating whether the symbol was chosen or rewarded, respectively, and *β* terms are their logistic regression coefficients with *β*_0_ representing the intercept. Model 5 includes their interaction. We defined *α* as a logistic function in order to constrain it neatly to between 0 and 1.

Model 6 then added fixation times to the specification for Model 5:

|  |  |
| --- | --- |
| Model 6 | $\alpha_t = \frac{1}{1 + e^{-(\beta_0 + \beta_{rewarded} r_t + \beta_{chosen} c_t + \beta_{interaction} r_t c_t + \beta_{fixation} f_t)}}$ |

where *f*_*t*_ ∈ [0, 1] is the portion of the feedback period spent fixating on the item, and *β*_*fixation*_ is its logistic regression coefficient.

Learning models were fit using the No-Uturn Sampling (NUTS) algorithm in Stan 2.9 (Stan Development Team, 2016), a Bayesian sampling method that estimates the most likely parameter values while providing a distribution of each value’s likelihood given the data. Models were estimated hierarchically, meaning that each participant had their own parameter value that was assumed to result from a group distribution defined by hyperparameters. This served to constrain participant-level outliers and provide group mean estimates for each parameter, an indicator of the likely generative value in the sample. We estimated the predictiveness of each model using a leave-one-out cross-validation procedure, which generates an expected log pointwise predictive density (ELPD) for data from a new experiment, as well as a standard error for each ELPD estimate. ELPDs were estimated and compared using the LOO package for R (Vehtari et al., 2015)

